# Broadly neutralizing antibody-secreting CAR-T cells elicit Fc-mediated effector functions *in vitro* and suppress HIV in humanized mice

**DOI:** 10.64898/2026.03.10.710742

**Authors:** Zoe Stylianidou, Sarah Gerlo, Magdalena Wejda, Elianne Burg, Evelien De Smet, Ytse Noppe, Maxime Verschoore, Jolien Van Cleemput, Linos Vandekerckhove, Wojciech Witkowski

**Author notes:** **Correspondence:** Linos Vandekerckhove.

## Abstract

Despite significant advances in antiretroviral therapy (ART) that have transformed human immunodeficiency virus (HIV) infection from a fatal diagnosis to a manageable chronic condition, the persistent viral reservoir necessitates lifelong treatment underscoring the critical need for curative interventions. Viral persistence within anatomically distinct reservoirs, coupled with HIV-associated immune dysregulation accentuates the need for innovative combination immunotherapies that can act through multiple mechanisms. We introduce the Hybrid chimeric antigen receptor (CAR) platform: a dual-function immunotherapy that combines the targeted cytotoxicity of CAR-T cells with the secretion of broadly neutralizing antibodies (bNAbs). This approach enables direct elimination of HIV-infected cells, neutralization of free virus and Fc-mediated effector recruitment. In vitro, Hybrid CAR-T cells eliminated HIV-infected CD4+ T cells while the secreted bNAbs neutralized HIV and mediated robust Fc-effector functions including antibody-dependent cellular cytotoxicity (ADCC) and antibody-dependent cellular phagocytosis (ADCP). In humanized mice, Hybrid CAR-T treatment achieved more than a 9-fold reduction in plasma viremia, accompanied by a significant decrease in viral levels across tissues, with circulating bNAbs detected in plasma. Collectively, these findings highlight the potential of Hybrid CAR-T cells as a synergistic next-generation therapeutic that bridges cellular and humoral immunity, supporting their translational promise as a strategy toward functional HIV cure.

## Introduction

Human immunodeficiency virus (HIV) remains a major public health issue, affecting 40.8 million adults and children globally (*1*). While antiretroviral therapy (ART) suppresses the virus and improves life expectancy of people living with HIV (PLWH), it is not curative (*2*). HIV forms stable latent reservoirs during early infection and ART interruption leads to viral rebound, necessitating lifelong treatment (*3–6*). However, long-term ART administration is associated with cumulative toxicities, increased risk of comorbidities and substantial financial burden (*7–9*). Beyond physiological consequences, the emotional and psychological burden experienced by PLWH remains significant, as HIV-related stigma and discrimination continue to negatively impact medication adherence and mental health (*10*). Together these challenges underscore the urgent need for curative strategies (*11*). To date, ten individuals have achieved either a cure or long-term remission of HIV, following hematopoietic stem cell transplantation to treat underlying hematologic malignancies; an approach that carries substantial risks and lacks scalability for broader clinical implementation (*12–20*). Consequently, more scalable, less invasive therapy options are a central research focus.

Cell and gene therapies have emerged as HIV cure candidates, addressing barriers like antigen escape and immune evasion, offering long-term immune surveillance and holding potential for a functional cure (*21–24*). Chimeric antigen receptor (CAR)-T cells, which have revolutionized the treatment of hematologic malignancies, reprogram T cells for Major Histocompatibility Complex (MHC)-independent antigen recognition and trigger potent cytotoxic responses via CD3ζ and costimulatory domains (*25–30*). Building on these successes, CAR-T therapy for HIV has advanced through successive design refinements. First generation CARs demonstrated a safety profile but failed to achieve sustained viral control, while later designs incorporating costimulatory domains improved T cell functionality and persistence (*31–34*). Clinical evaluation of broadly neutralizing antibody (bNAb)-derived, third-generation CAR-T cells showed delayed viral rebound following treatment interruption, although viral escape remained a challenge, while current clinical trials continue to assess CAR-T cell safety and efficacy (*35–40*).

In parallel, direct administration of bNAbs has demonstrated both safety and antiviral activity; however when used as monotherapy provides incomplete viral control (*41,42*). Consequently, emerging combination strategies, particularly those incorporating cell-based approaches, aim to enhance reservoir clearance and promote durable remission (*43,44*). A key feature of bNAbs is the IgG1-Fc domain which facilitates engagement with Fcγ receptors (FcγR) on innate immune cells, thereby recruiting effector mechanisms such as antibody-dependent cellular cytotoxicity (ADCC) and antibody-dependent cellular phagocytosis (ADCP), both of which have been associated with antiviral activity (*45,46*).

We introduce the Hybrid CAR platform, a dual-function therapeutic targeting both HIV-infected cells and free virus. Hybrid CAR-T cells co-express a CD4-based anti-HIV CAR and secrete the 3BNC117 single chain variable fragment fused to human IgG1-Fc (3BNC117scFv-IgG1Fc). In this bicistronic design, the CD4-based CAR mediates MHC-independent cytotoxicity, while the secreted 3BNC117scFv-IgG1Fc neutralizes cell free virus and engages Fc-dependent effector functions, including ADCC and ADCP, thereby integrating cellular and humoral immune responses into a single product.

## Materials and methods

### Study design

The goal of this study was to preclinically evaluate the functional activity and antiviral potential of the Hybrid CAR-T cell platform which combines the CD4-based CAR-driven direct cytotoxicity with the secretion of a bNAb-based molecule (3BNC117scFv-IgG1Fc). A bicistronic lentiviral vector was constructed to co-express the CAR and the antibody via a T2A self-cleaving peptide, allowing coordinated production from a single transcript. Primary human CD8+ T cells isolated from HIV-negative donors were transduced and assessed *in vitro* for CAR expression, cytotoxicity against HIV Env-expressing targets, antibody secretion, neutralization of HIV and Fc-mediated effector functions including ADCC and ADCP. For *in vivo* evaluation, PBMC-humanized NSG-SGM3 mice were engrafted with human immune cells and infused with infected CD4+ T cells, followed by autologous Hybrid CAR-T cell administration. Key endpoints included plasma viremia, tissue-associated DNA and circulating antibody levels quantification.

### Plasmids

The lentiviral transfer vector encoding an anti-HIV CAR with a CD4 extracellular domain, CD8a hinge, CD28 transmembrane and intracellular domain and 4-1BB/CD3ζ costimulatory domains was kindly provided by Prof. James L. Riley (*47*). The sequence encoding a self-cleaving T2A peptide, an interleukin-2 (IL-2) signal sequence and a 3BNC117-scFv fused to a human IgG1-Fc (hinge-CH2-CH3) was synthesized (Integrated DNA Technologies) and cloned downstream of CD3ζ using the Gibson Assembly cloning kit (New England Biolabs) (*48*). The CD4 CAR, Hybrid CAR and GFP-3BNC117 amino acid sequences are shown in Table S1. The proviral plasmid NLENG1-IRES engineered to express eGFP, the VSVg envelope and HIV-1 gag/pol/rev packaging were kindly provided by Prof. Bruno Verhasselt (Ghent University, Belgium) (*49*). The 3BNC117-resistant envelope x2088.c9 plasmid was kindly provided by Prof. Michael A Seaman and Dr. Christy Lavine (Beth Israel Deaconess Medical Center, United States) (*50*).

### Virus production and titration

For lentivirus production, Lenti-X 293 T cells (3.7×10^6^/10cm dish) were co-transfected 24h post-seeding with VSVg envelope (0.72pmol), HIV-1 gag/pol/rev packaging (1.3pmol) and transfer vector plasmid (1.64pmol) using Lipofectamine 2000 (Invitrogen). Medium was changed to DMEM (Gibco) supplemented with 10% FCS (Pan-Biotech) and 4mM L-glutamine (Gibco) at 16h. Supernatants were harvested at 72h, filtered (0.45μm), concentrated 10-fold (Amicon Ultra-15 100K) and stored at -80°C. Functional titers were determined by transducing Lenti-X 293T cells with serial dilutions in polybrene 8μg/ml (Merck Millipore) using spinoculation (1000xg, 90min, 32°C). Transduction efficiency was quantified by CD4 staining and flow cytometry (MACSQuant 10). Transducing units per millilitre (TU/mL) were calculated from the fraction of CD4+ cells and inoculum volume. Replication competent HIV_NL4.3-eGFP_ was produced by transfecting HEK293T cells with pNLENG1-IRES. HIV_X2088.c9_ pseudovirus was generated by co-transfecting x2088.c9 envelope with pSG3ΔEnv (NIH HIV Reagent Program, ARP-11051) (*51*). Supernatants were harvested at 48h, filtered (0.45μm), and stored at -80°C. Titers were determined using QuickTiter™ Lentivirus Titer Kit.

### PBMC isolation and cell culture

PBMCs were isolated from buffycoats using Lymphoprep density gradient centrifugation. CD4+, CD8+, CD14+ and NK cells were enriched by magnetic selection (Miltenyi Biotec) and cultured at 10^6^cells/ml. CD4+ T cells were maintained in TexMACS medium with 5% human serum (CELLect, MP Biomedicals), 1% penicillin-streptomycin (Gibco) and 20ng/ml IL-2 (PeproTech) and stimulated with anti-CD3/CD28 TransAct (Miltenyi Biotec) for 72h before infection. CD8+ T cells were cultured in TexMACS medium with 1% penicillin-streptomycin, 40IU/μl IL-7 (Miltenyi Biotec), 40IU/μl IL-15 (Miltenyi Biotec) and Transact. 24h post isolation, cells were transduced with lentiviral stock (MOI 25) in presence of polybrene 8μg/ml using spinoculation (1000xg, 90min, 32°C). On day 3, TransAct was removed and cells were expanded for 8 days in IL-7/IL-15 medium. CD14+ monocytes were cultured in RPMI (Gibco) with 10% FCS, 1% penicillin-streptomycin, 50ng/ml M-CSF (Miltentyi Biotec). Medium was refreshed on day 3 and cells differentiated for additional 4 days. NK cells were cultured in RPMI with 10% FCS, 1% penicillin-streptomycin, 10ng/ml IL-2 and 25IU/ml IL-15 for 24h before use.

### In vitro HIV suppression assays

Primary CD8+ T cells were transduced with Hybrid CAR lentivirus or left non-transduced (NTD). On day 7 post-transduction, effector cells were co-cultured with GFP+ Raji-Env target cells at 2:1 effector-to-target (E:T) ratio in RPMI with 5% human serum, 1% penicillin-streptomycin and 20ng/ml IL-2 for 48h at 37°C. Cells were stained with live/dead dye, CD8-APC (Miltenyi Biotec), CD4-APCVio770 (Miltneyi Biotec) and analyzed by flow cytometry (MACSQuant 10). Activated CD4+ T cells were infected with HIV_NL4.3-eGFP_ (MOI 1.25) for 4 days, then co-cultured with autologous CAR or NTD CD8+ T cells at E:T of 5:1, 1:1, 1:10. HIV replication was quantified by GFP+ events in the CD8negative population at days 2, 5 and 7. Specific lysis (%) was calculated as: 100-[(% GFP+ cells in Hybrid CAR ÷ % GFP+ cells in NTD) x 100]. Cytolysis kinetics were monitored continuously using the IncuCyte S3 Live-Cell Imaging System (Sartorius).

### TZM-bl assay

Infectivity of co-culture supernatants was assessed using the TZM-bl reporter cells. Co-culture supernatants were mixed with TZM-bl cells at 37°C. After 48h, cells were washed and treated with β-galactosidase substrate (Pierce™, Thermo Scientific) and absorbance was measured at 405nm. For neutralization assays, supernatants from NTD or Hybrid CAR CD8⁺ T cell cultures were mixed with pre-titrated HIV_NL4.3-GFP_ virus and incubated for 30min at 37°C. HIV_x2088.c9_ served as specificity control. Virus-supernatant mixtures were added to TZM-bl cells and processed as above. Virus infectivity was normalized to the ‘no antibody’ condition (100%) and neutralization (%) was calculated as: %Neutralization= 100-%Virus infectivity.

### Intracellular cytokine staining

Non-transduced or (Hybrid) CAR-transduced CD8+ T cells were cocultured with wild-type or HIV-1 YU-2 gp160-expressing Raji cells at 1:1 E:T ratio for 16-18h. Then, brefeldin A and monensin was added to the cocultures for 6h to accumulate the cytokines produced intracellularly. At the end of the coculture, cells were permeabilized and stained with anti-human CD8-APC (Miltenyi Biotec), CD4-APCVio770 (Miltneyi Biotec), ΤΝFα-Pacific Blue (Biolegend) and IFNγ-PE (Miltenyi Biotec) and analyzed by flow cytometry (MACSQuant 10).

### CAR-T cell-secreted bNAb 3BNC117 quantification by Enzyme Linked Immunosorbent Assay (ELISA)

MaxiSorp 96-well plates (Thermo Scientific) were coated overnight at 4°C with 200ng/ml recombinant HIV-1 gp120 protein (Sino Biological). Wells were washed with PBS-0,1% Tween20 and blocked with B3T buffer (150mM NaCl, 50mM Tris-HCl, 1mM EDTA,3.3% FCS, 2%BSA, 0.07% Tween20). Cell culture supernatants and serial dilutions of recombinant 3BNC117 (NIH HIV Reagent Program, ARP-12474) (*52*). After washing, HRP-conjugated anti-human IgG-Fc (Bethyl Laboratories) was added for 1h at 37°C, wells were washed, TMB substrate (Merck Millipore) was added for 15min at RT, reaction was stopped with 1M H2SO4 and absorbance was measured at 450nm.

### Antibody-dependent natural killer cell activation and degranulation (ADNKDA)

ADNKDA assay was performed as described elsewhere (*53*). Briefly, HIV-1 gp120 protein was coated onto MaxiSorp plates at 500ng/well at 4°C overnight. Wells were washed with PBS, blocked with PBS-5% BSA, then incubated with cell supernatants or serial dilutions of recombinant 3BNC117 for 2h at 37°C. After washing, NK cells (5×10^4^/well) were added with protein transport inhibitor (Invitrogen) and anti-CD107a (Miltenyi Biotec) for 5h at 37°C. Cells were analyzed with flow cytometry (MACSQuant 10).

### Antibody-dependent cellular phagocytosis (ADCP) with monocyte-derived macrophages

CD14+ monocyte-derived macrophages were seeded on a flat-bottom 96-well plate (5×10^4^/well), in RPMI 1640 with GlutaMAX with 10% FBS and 1% penicillin-streptomycin. HIV-1 YU-2 gp160-expressing Raji target cells were labelled with IncuCyte pHrodo Red Dye (Sartorius) and incubated with bNAb-containing cell culture supernatants. Serial 2-fold dilutions of recombinant 3BNC117 served as controls. Opsonized targets were added to macrophages and phagocytosis was tracked live using the IncuCyte instrument. Fluorescence was quantified as ‘Total Red Object Integrated Intensity’.

### Humanized Mice experiments

6-8 week old NSG-SGM3 (NOD.Cg-Prkdcscid Il2rgtm1Wjl Tg(CMV-IL3,CSF2,KITLG)1Eav/ MloySzJ) mice (JAX, Stock No. 013062) of both sexes, used at equal frequences, were retro-orbitally injected with 10×10^6^ CD19-depleted PBMCs derived from HIV-negative donors. Twelve days later, mice received 10×10^6^ autologous HIV_NL4.3_-infected CD4+ T cells and combination ART (dolutegravir/abacavir/lamivudine; Triumeq®, ViiV Healthcare) for 4 days. Crushed tablets were dissolved in Sucralose MediDrop® (Clear H2O) and incorporated into DietGel® Boost cups (Clear H2O) at the dose of 20mg per cup. On day 16, mice received 5×10^6^ autologous non- or CAR-transduced CD8+ T cells. All cell suspensions were resuspended in 200μl Hanks’ Balanced Salt Solution (HBSS, Gibco) before administration. Peripheral blood was collected via retro-orbital bleeding and stained with anti-murine CD45 PE-Cy7 (Biolegend), anti-human CD45 Pacific Blue (Biolegend), anti-human CD3 PE-Cy5.5 (Invitrogen), anti-human CD4 PE (BD Biosciences) and anti-human CD8 (BD Biosciences) for flow cytometry (BD LSR Fortessa), to assess engraftment. At the study endpoint, spleen, lung and bone marrow tissues were harvested; half of were stored at -80°C for quantification of viral copies half processed into single-cell suspensions for phenotyping. Mice showing severe graft versus host disease (GvHD) were humanely euthanized. Mouse experiments were performed following a protocol adapted from Leibman et al (*47*).

### HIV RNA viral load assay

HIV RNA was isolated from 50μl of plasma using the QIAamp Viral RNA Mini Kit (Qiagen) and reverse transcribed using Random Hexamers (Invitrogen) and SuperScript™ IV Reverse Transcriptase (Invitrogen). Viral loads were determined by qPCR using the LightCycler® 480 Probes Master (Roche) on the LightCycler® 480 System. The *int* gene was targeted with the following primer/probe set: Forward primer: 5’-GGTTTATTACAGGGACAGCAGAGA-3’, Reverse primer: 5’-ACCTGCCATCTGTTTTCCATA-3’, Probe: 5’-FAM-ACTACTGCCCCTTCA CCTTTCCARAG-BHQ1-3’. A 10-fold dilution series of the pNLENG1-IRES plasmid was used to generate the standard curve.

### HIV DNA quantification in murine tissues

gDNA from the spleen, lung and bone marrow was isolated using the DNeasy Blood & Tissue Kit (Qiagen). Total HIV-1 DNA was quantified by digital PCR on the QIAcuity System (Qiagen) with ≥500ng gDNA per reaction. The 40μl reaction contained 10μl 4x Qiacuity Probe Master Mix (Qiagen), 2μl *int* primer/probe set, 5μl template gDNA and nuclease free water. Reactions were run in duplicate on 26K 24-well nanoplates (Qiagen) and in the following conditions: an initial activation step at 95°C for 2 minutes, followed by 45 cycles at 95°C for 30 seconds and 58°C for 60 seconds. 5μl of 1:50 diluted gDNA was used for the RPP30 reference gene quantification, as described elsewhere (*54*).

### CAR distribution in murine tissues

Digital PCR was performed to assess CAR-T cell distribution in murine tissue. CAR transgene abundance was quantified using the following primer/probe set: Forward primer: 5’-AAG CAT TAC CAG CCC TAT GC -3’, Reverse primer: 5’-AGT TCA CAT CCT CCT TCT TC -3’, Probe: 5’-FAM - TAC AAA CTA CTC AAG AGG AAG ATG GCT GT – BHQ1 – 3’. GFP-3BNC117 transgene abundance was quantified using the following primer/probe set: 5’-AAT TGT TAC AGT CTG GGG CAG -3’, Reverse primer: 5’-AAT GTG TCA AAG TCC CAC G -3’, Probe: 5’-HEX - CAA TCC TAA GAC AGG TCA GCC AAA CAA TC -3’.

### Ethics statement

Peripheral blood mononuclear cells (PBMCs) from HIV-negative donors were obtained from the Biobank Red Cross-Flanders Center after informed consent (Biobank Red Cross-Flanders, G20240422A) and with approval from the Ethical Committee of Ghent University (BC-10607). All in vivo mouse experiments were conducted according to the Federation of European Laboratory Animal Science (FELASA) guidelines and approved by the Ethical Committee of Ghent University (ECD 21/33K, ECD 22/65).

### Statistical analysis

All statistical analysis was performed using GraphPad Prism (version 10.5.0, GraphPad Software, San Diego, CA, USA). For paired comparisons between supernatants derived from the same donor, the non-parametric Wilcoxon signed-rank test was applied. For comparisons involving more than two independent groups, the non-parametric Kruskal-Wallis test was used. In figures, significance levels are denoted as: ns = non-significant, *p<0.05, **p<0.01, ***p<0.001, ****p<0.0001.

## Results

### Design of the Hybrid CAR construct and genetic engineering of anti-HIV Hybrid CAR-T cells

We utilized a previously described CD4-based CAR targeting the HIV envelope. This CAR construct incorporates a CD8a hinge and transmembrane domain, followed by CD28 and 4-1BB costimulatory domains and a CD3ζ activation domain, enabling robust T cell activation and persistence (47). Based on this construct, we generated two bicistronic lentiviral vectors: a vector designed to co-express two synergistic anti-HIV modalities: (i) the anti-HIV CD4-based CAR and (ii) a secreted 3BNC117 single-chain variable fragment fused to human IgG1-Fc (3BNC117scFv-IgG1Fc), and a control bicistronic vector in which the CD4 CAR is replaced with GFP (table S1). A T2A self-cleaving peptide sequence is positioned between the CAR and the 3BNC117scFv-IgG1-Fc ensuring separate expression of both molecules (Fig. 1A). Flow cytometry was used to evaluate transduction efficiency by measuring the level of CD4 surface expression in primary CD8+ T cells from HIV-negative donors. Gating thresholds were set on non-transduced cells (NTD), with CAR+ cells identified as CD4+CD8+ double-positive events and expression was confirmed across donors (11.8 ± 7.1%; Fig. 1, B and C, fig. S1).

**Fig. 1.**
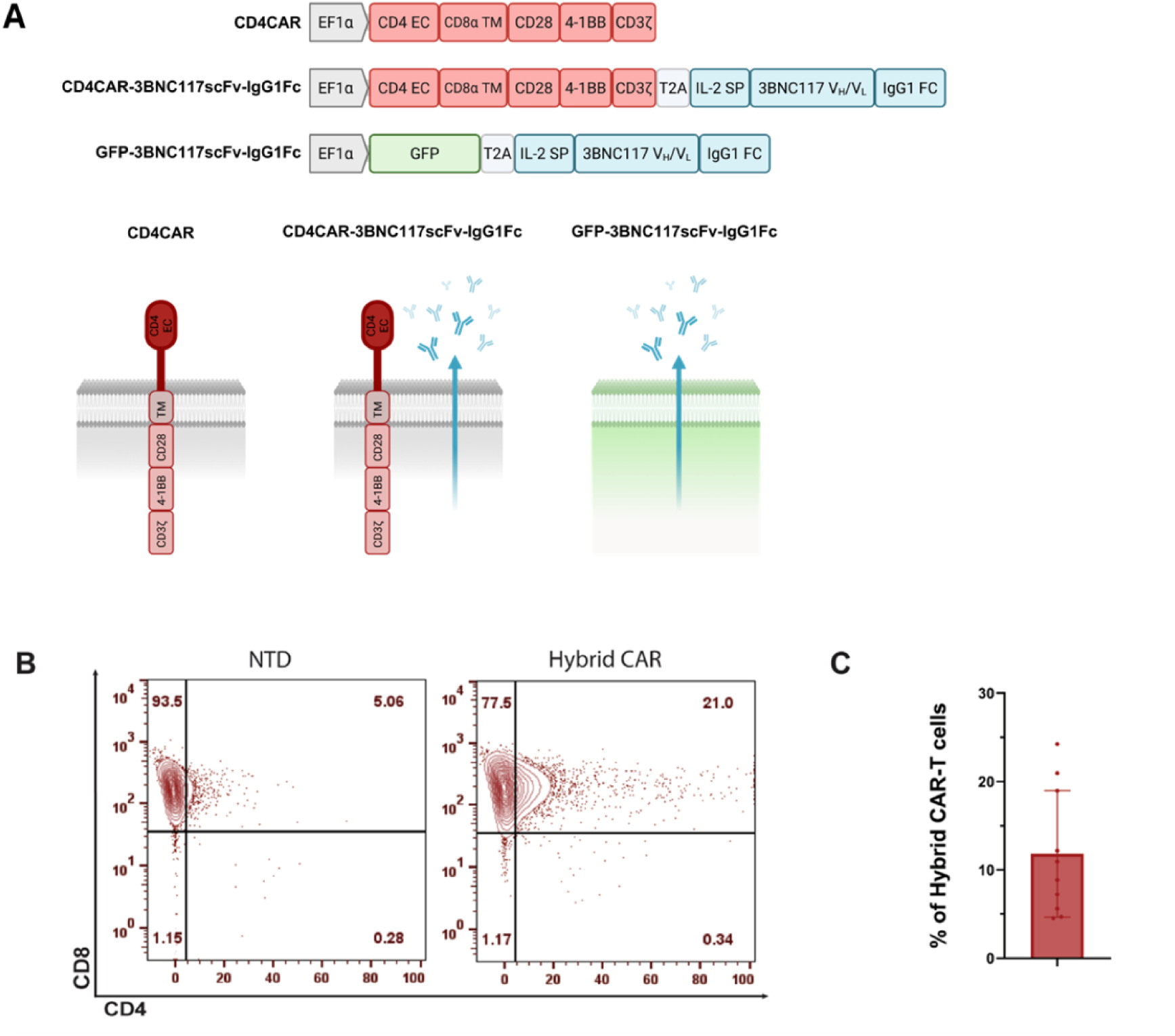
Generation of Hybrid CAR-T cells. **(A)** Schematic representation of lentiviral transfer vectors used in this study. Hybrid CAR-T cells were generated via lentiviral transduction using a bicistronic vector. Expression of the encoded fusion protein is under the control of the EF1α promoter. The CD4 CAR and the 3BNC117scFv-IgG1Fc coding regions are separated by a T2A cleavage sequence. Control vectors include one encoding the CD4 CAR alone and a vector encoding GFP fused to 3BNC117scFv-IgG1Fc. **(B)** Representative flow plot showing surface CAR expression transduced primary CD8+ T cells. CAR detection was measured by gating for CD4+CD8+ T cells, representing the Hybrid CAR-transduced cells. **(C)** Graphical representation of % Hybrid CAR-modified T cells from multiple donor. Each dot represents one donor; n=10. Abbreviations: CD4 EC, CD4 extracellular domain; CD8α TM, CD8α transmembrane domain; IL-2 SP, IL-2 signaling peptide; NTD, non-transduced.

### Hybrid CAR-T cells exert strong cytotoxic effects against Env-expressing targets and autologous HIV-infected CD4+ T cells in vitro

To assess their antiviral activity, Hybrid CAR-T cells derived from primary CD8+ T cells were co-cultured with Raji cells stably expressing HIV-1 YU-2 Env, at 2:1 effector-to-target (E:T) ratio for 48h. Flow cytometry revealed specific lysis of Env-expressing targets by Hybrid CAR-T cells across donors (Fig. 2A). Next, we compared the cytotoxic activity of Hybrid CAR-T cells to conventional CD4 CAR-T cells and appropriate controls against autologous primary CD4+ T cells infected with replication competent HIVNL4.3-eGFP. Non-transduced (NTD) and GFP-3BNC117-transduced T cells, which secrete bNAbs but lack CAR expression, were included as negative and antibody-only controls, respectively. Cells were co-cultured with HIV-infected targets at E:T ratios of 5:1, 1:1 and 1:10 for 7 days. HIV infection dynamics were quantified by flow cytometry as GFP+ infected cells on days 2, 5 and 7 (fig. S2). Supernatants were also collected at the same time points and assessed using a TZM-bl assay to determine levels of infectious virus. In parallel, continuous live-cell imaging was performed to monitor infection dynamics and CAR-T cell activity in real time.

**Fig. 2.**
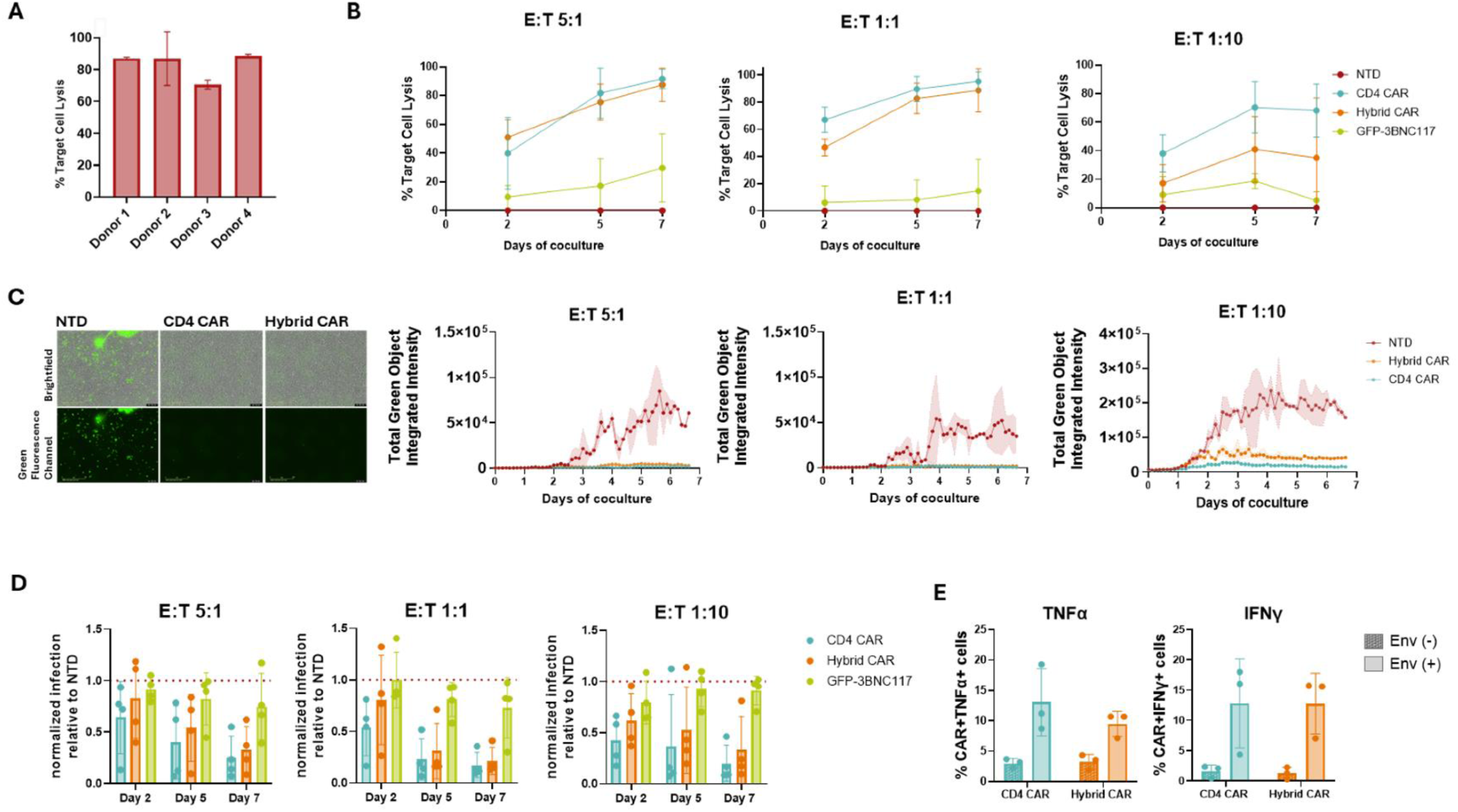
Anti-HIV Hybrid CAR-T cells exert HIV-specific killing in vitro. **(A)** Cytotoxic assay of Hybrid CAR-T cells co-cultured with Env-expressing Raji cells at E:T 2:1 for 48h; n=4. **(B)** Cytotoxic assay of Hybrid CAR-T cells co-cultured with autologous HIV_NL4.3_-infected CD4+ T cells at E:T 5:1, 1:1 and 1:10 for 7 days; each dot represents the mean percentage target lysis from four donors; n=4. **(C)** (left) Representative live-cell images in Brightfield and in green fluorescence channel captured by the IncuCyte system, at E:T 5:1, on day 7 of coculture. Images were captured at 20x magnification. (right) Representative kinetic curves showing HIV replication in real time over the 7-day observation period of the coculture. **(D)** TZM-bl assay measuring HIV infectivity in supernatants collected at day 2, 5 and 7 of coculture. Data were normalized to the NTD condition, which was set to 1. The dotted line indicates the normalized NTD level; Each dot represents one donor; n=4. **(E)** Percentage of cytokine-producing CAR-T cells in presence of Env+ targets; each dot represents one donor; n=3. Abbreviations: NTD, non-transduced.

At higher E:T rations (5:1 and 1:1), Hybrid CAR-T cells exhibited cytotoxic activity comparable to CD4 CAR-T cells across all time points. Under more stringent conditions with limiting effector cell numbers (E:T 1:10) and in a subset of donors, Hybrid CAR-T cells showed a modest reduction in killing efficiency relative to CD4 CAR-T cells, while maintaining significantly greater cytotoxic activity than control conditions (Fig. 2B). Consistent with these findings, live-cell imaging confirmed sustained control of infection in CD4 CAR-T and Hybrid CAR-T cocultures over the 7-day period (Fig. 2C, supplementary videos 1-3). TZM-bl reporter cell assay revealed a reduction of infectious virus particles in supernatants from both CAR conditions, with only partial suppression observed for GFP-3BNC117-transduced T cells (Fig. 2D). As shown in Fig. 2E, co-culture of CD4 CAR-T and Hybrid CAR-T cells with Env-expressing target cells triggered TNFα and IFNγ production, whereas minimal cytokine secretion was detected in the presence of Env-negative targets. In addition, CAR-transduced cells exhibited antigen-specific activation, as evidenced by upregulation of the CD69 activation marker and expansion during co-culture, whereas GFP-3BNC117-transduced T cells showed no detectable activation or expansion (fig. S3 and S4). Lastly, to evaluate whether the extracellular CD4 moiety of the Hybrid CAR could serve as an entry receptor for HIV, CD4 knock-out cells were transduced with the Hybrid CAR construct and subsequently exposed to HIV. Despite efficient CAR expression, no productive infection was detected, indicating that the CD4 domain in the CAR does not render the modified T cells susceptible to HIV infection (fig. S5).

### Hybrid CAR-T cells secrete functional 3BNC117scFv-IgG1Fc, conferring HIV-neutralizing activity

Five days post-transduction, culture supernatants of CAR-transduced, GFP-3BNC117-transduced and non-transduced CD8+ T cells, were harvested and analyzed for the presence of the secreted 3BNC117scFv-IgG1Fc antibody. Quantification by ELISA confirmed consistent expression of the antibody from Hybrid CAR-T cells across donors, with a mean concentration of 34.91 ng/mL (range: 19.59-54.11 ng/mL; n=8) (Fig. 3A). Due to higher transduction efficiencies, increased antibody levels were observed in GFP-3BNC117-transduced T cells. To assess whether the secreted antibody retained functional antiviral activity, neutralization capacity was evaluated using the TZM-bl reporter cell assay. Infection levels were normalized to the ‘no antibody’ control condition. Supernatants derived from Hybrid CAR-T cells achieved a mean neutralization of 41.13% (41.13 ± 12.38%), indicating that the secreted antibody maintained robust neutralizing functionality following expression and processing in T cells. GFP-3BNC117 T cell-supernatants showed higher neutralization consistent with elevated antibody levels. Non-transduced and CD4 CAR-T supernatants exhibited negligible inhibitory activity, confirming that neutralization was specifically attributable to secreted antibodies. As an additional specificity control, neutralization assays were also performed against the 3BNC117-resistant HIV-1 isolate x2088.c9. Under these conditions, Hybrid CAR- and GFP-3BNC117 T cell supernatants showed no measurable to minimal inhibition, validating strain-specific neutralizing activity (Fig. 3B, fig. S6). Collectively, these results demonstrate that Hybrid CAR-T cells not only express the CAR construct but also successfully secrete fully functional antibodies capable of mediating antiviral activity.

**Fig. 3.**
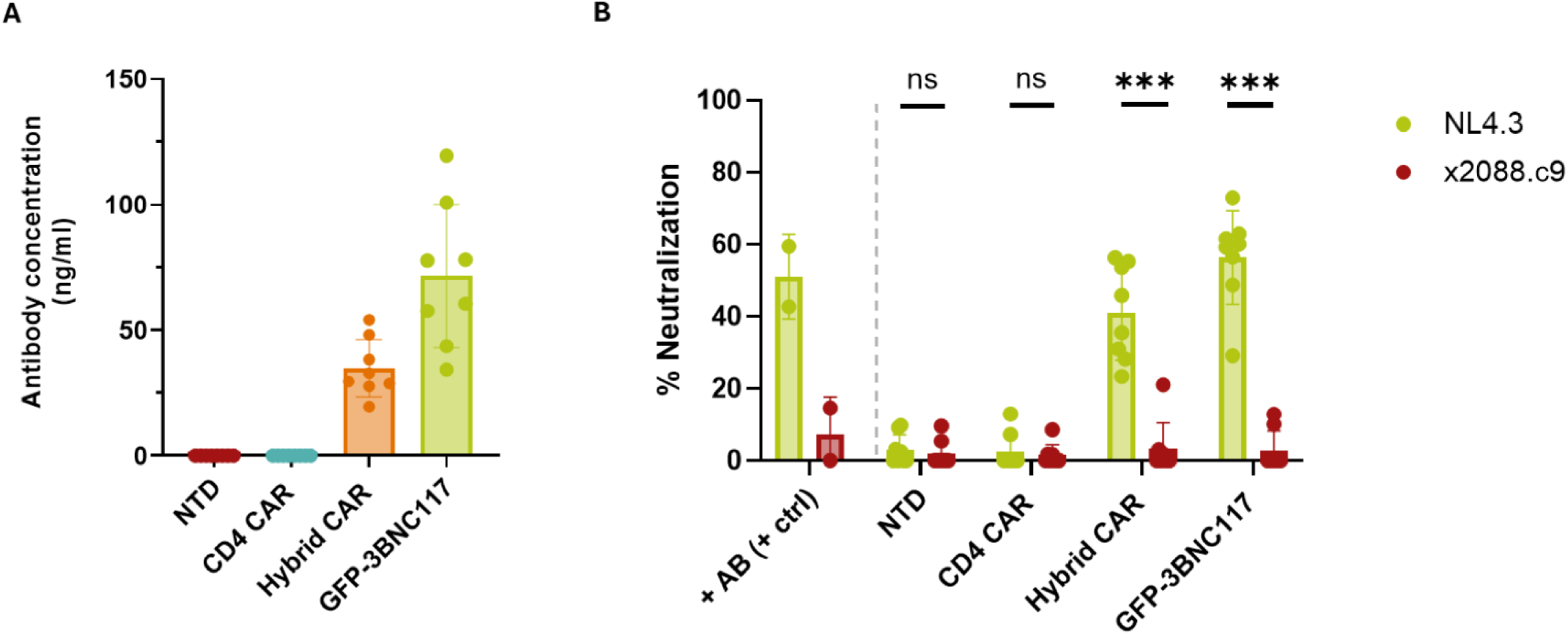
Hybrid CAR-T cells secrete 3BNC117scFv-IgG1Fc with HIV-neutralizing capacity. **(A)** Mean antibody concentration of 3BNC117scFv-IgG1Fc in the supernatants of cells transduced with the indicated lentiviral constructs, as measured by ELISA; n=8. **(B)** Cell culture supernatants can neutralize HIV_NL4.3_ but not HIV_x088.c9_ (Wilcoxon signed-rank test; p values indicated with ns = non-significant, ***p<0.001; Each dot represents one donor; n=8. Abbreviations: +AB (+ctrl), recombinant antibody (positive) control; NTD, non-transduced.

### Hybrid CAR-T cell-secreted antibodies mediate Fc-effector functions in vitro

Because Fc-mediated effector functions are known to contribute to HIV immunity, we investigated whether antibodies secreted by Hybrid CAR-T cells are capable of driving both ADCC and ADCP. ADCC activity was evaluated using an antibody-dependent NK cell degranulation assay (ADNKDA) which measures the activation of primary NK cells in response to antibody-coated targets by assessing surface expression of the degranulation marker CD107a. Particularly, supernatants derived from Hybrid CAR-T cells significantly enhanced primary NK cell degranulation by 194.33% compared to supernatants from non-transduced cells (mean 194.33 ± 76.92%; p=0.0039; n=9), indicating that the secreted 3BNC117scFv-IgG1Fc antibodies effectively engage NK cells and trigger cytotoxic effector functions (Fig. 4A). ADCP was evaluated in real time using Env-expressing Raji target cells, labelled with a pH-sensitive dye that fluoresces red upon exposure to the acidic environment of phagolysosomes and allowing direct visualization of antibody-mediated phagocytic uptake. Signal from the phagocytosed target cells reached maximum at approximately 60-90 minutes before gradually declining over the subsequent 4-hour observation (Fig. 4B). At the 1-hour timepoint, supernatants from Hybrid CAR-T cells induced a 3.33-fold increase in red fluorescence compared to non-transduced controls (3.33 ± 1.16 SD; p=0.0312; n=6) (Fig. 4C). Importantly, monocyte-derived macrophages exhibited strong red fluorescence exclusively in the presence of 3BNC117scFv-IgG1Fc-containing supernatants, consistent with Fc-dependent uptake and confirming the functional engagement of innate phagocytic mechanisms (Fig. 4D). In both assays, supernatants from GFP-3BNC117-transduced cells induced higher effector functions, consistent with their elevated antibody levels. Together, these data suggest that antibodies secreted by Hybrid CAR-T cells not only retain neutralizing activity but are also capable of recruiting and activating innate immune effector cells, mediating NK cell cytotoxicity via CD107a degranulation and macrophage phagocytosis, thereby enhancing the potential antiviral activity of this dual-function therapeutic approach.

**Fig. 4.**
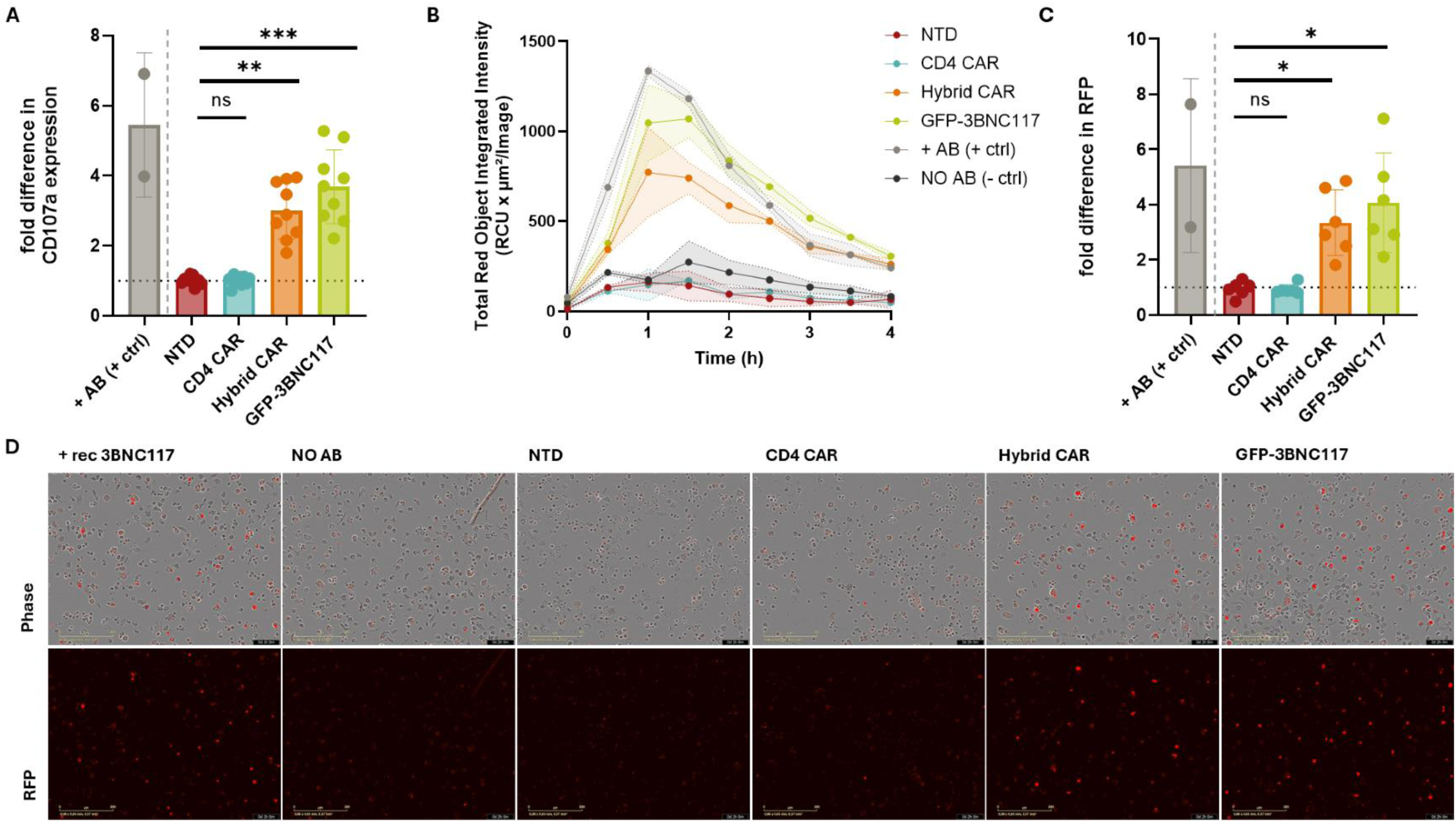
Hybrid CAR-T cell-secreted 3BNC117scFv-IgG1Fc elicit Fc-mediated effector functions. **(A)** Primary human NK cells were analyzed for surface expression of the degranulation marker CD107a (One-way ANOVA, Kruskal-Wallis test; p values indicated with **p<0.01, ***p<0.001; n=9). **(B)** Representative kinetic curves showing the amount of red fluorescence emitted over the 4h observation period of the ADCP assay. **(C)** Amount of red fluorescence emitted at the peak of the ADCP assay (One-way ANOVA, Kruskal-Wallis test; p values indicated with *p<0.05; Each dot represents one donor; n=6). **(D)** Representative live-cell images in Brightfield and in red fluorescence channel captured by the IncuCyte system, at the peak of the ADCP assay. Images were captured at 20x magnification. Data were normalized to the -AB condition, which was set to 1. The dotted line indicates the normalized - AB level. Abbreviations: -AB, no antibody (negative) control; +AB (+ctrl), recombinant antibody (positive) control; NTD, non-transduced.

### Hybrid CAR-T cells mediate potent suppression of HIV and systemic antibody delivery in vivo

After confirming the cytotoxic effectiveness of Hybrid CAR-T cells in vitro, we next evaluated their performance in vivo. To optimize the in vivo model, we initially compared human immune cell reconstitution in NOD.Cg-Prkdcscid-Il2rgtm1Wjl/SzJ (NSG) and NOD.Cg-Prkdcscid-Il2rgtm1WjlTg(CMV-IL3,CSF2,KITLG)1Eav/MloySzJ (NSG-SGM3) mice. NSG-SGM3 mice demonstrated faster and more robust humanization and were therefore selected for subsequent studies (fig. S7). Successful humanization was confirmed in all animals at 4 weeks post-engraftment with human CD45+ cell frequencies reaching up to 42,6% in peripheral blood, 81,5% in the spleen, 73,3% in the lung and 6% in the bone marrow (fig. S8). Because graft versus host disease (GvHD) typically emerges between 3 to 5 weeks after humanization in this model, follow-up was limited to 4 weeks (Fig. 5A). On day 12 post-humanization, mice received autologous CD4+ T cells infected with HIVNL4.3, with concurrent ART initiation. To mitigate early onset of GvHD, CD4⁺ T cells were infected ex vivo before infusion. Four days later, Hybrid CAR T cells were administered and ART was subsequently interrupted to assess the in vivo antiviral activity of the CAR-T cells. As shown in Fig. 5B (left panel), baseline plasma viremia prior to CD8+ T cell infusion was comparable between groups (NTD: 0,229×105, CD4 CAR: 0,242×105, Hybrid CAR: 0,179×105 0,179×106 and GFP-3BNC117: 0,190×105 HIV RNA copies/ml of plasma; p>0,9999), confirming uniform infection levels. By 14 days post CAR-infusion, mice treated with Hybrid CAR-T cells exhibited a significant 9.38-fold reduction in plasma viremia compared to mice receiving non-transduced CD8+ T cells (p=0.0037) (Fig. 5B, right). In comparison, treatment with CD4 CAR-T cells resulted in a 5.35-fold reduction in plasma viremia, which did not reach statistical significance (p=0,0913), while mice receiving GFP-3BNC117 cells showed a modest 2,48-fold reduction (p=0,7846), indicating partial viral inhibition mediated by secreted bNAbs in the absence of CAR-dependent targeting.

**Fig. 5.**
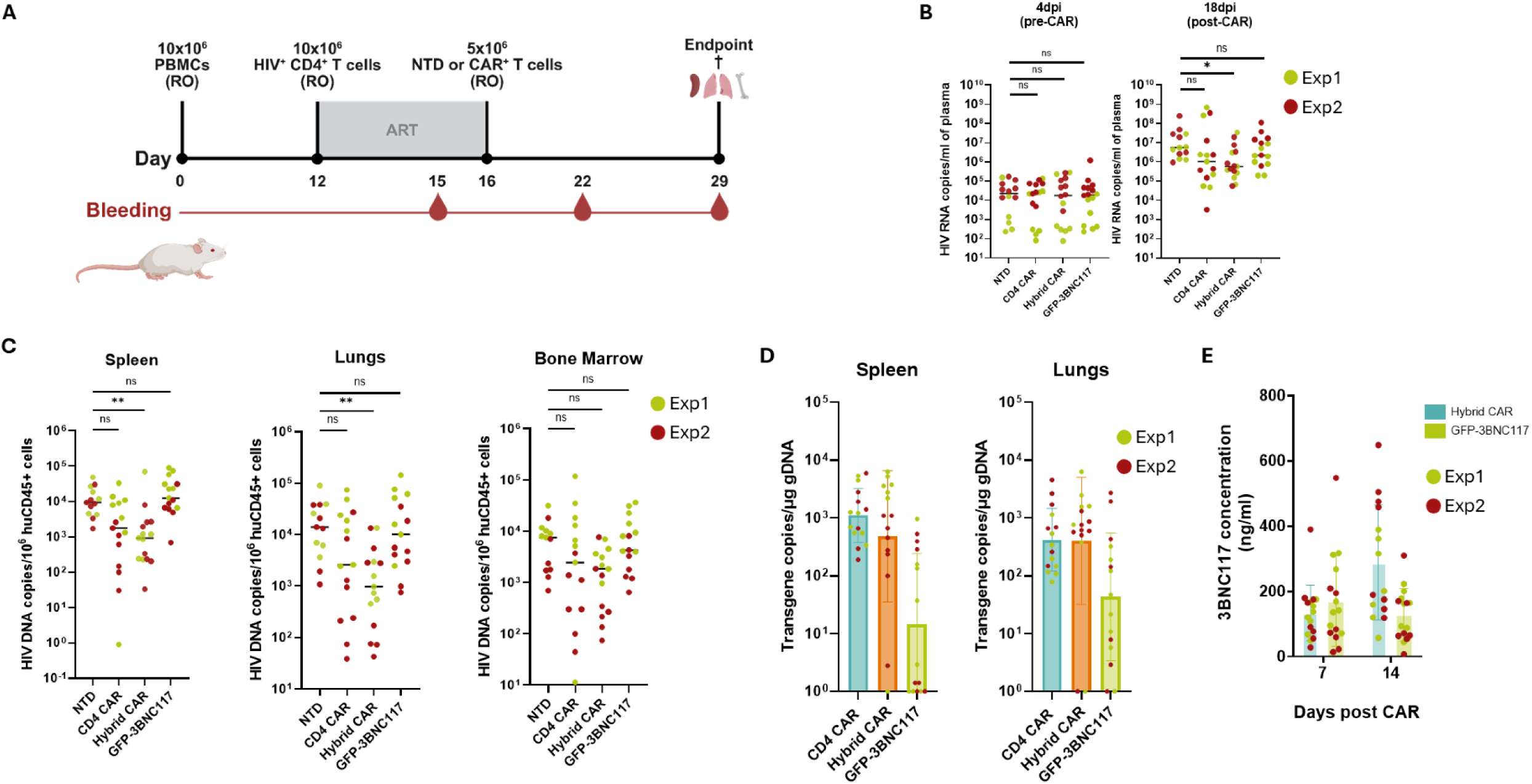
*In vivo* cytotoxic activity and sustained bNAb secretion by Hybrid CAR-T cells in humanized mice. **(A)** Experimental design of the in vivo experiment. **(B)** Plasma HIV viral loads of non-treated and Hybrid CAR-T cell treated mice expressed as copies per ml of plasma, as measured by RT-qPCR. Hybrid CAR-T cell-treated mice exhibited significantly lower viremia levels compared to non-treated mice. (One-way ANOVA, Kruskal-Wallis test; p values indicated with ns = non-significant, *p<0.05). **(C)** HIV-DNA levels in murine tissue compartments expressed as DNA copies per 106 huCD45+ cells, as measured by dPCR. Hybrid CAR-T cell-treated mice demonstrated significantly lower viral DNA levels in the spleen, the lungs and the bone marrow (One-way ANOVA, Kruskal-Wallis test; p values indicated with ns = non-significant, **p<0.01). **(D)** Plasma levels of 3BNC117scFv-IgG1Fc, 7 and 14 days post CAR administration as measured by ELISA. Each dot represents one animal. **(E)** CAR-T cell distribution in the spleen and the lungs. CAR gene copies were detected by dPCR; n=14, Exp1; n=16, Exp2. Abbreviations: RO; Retro-orbitally, NTD: non-transduced; Exp: Experiment. Data combined from two individual mouse experiments, each one indicated with a dedicated color.

Quantitative tissue analysis revealed significant reductions in HIV DNA levels across multiple anatomical compartments following Hybrid CAR treatment. Specifically, the spleen tissues displayed a 10.11-fold decrease (p=0.0087), lung tissues showed a 14.34-fold decrease (p=0.0002) and bone marrow HIV DNA levels were reduced 4.09-fold (p=0.0506) (Fig. 5C). CD4 CAR-T cell treatment was associated with more moderate, non-significant reductions in tissue-associated HIV DNA (5,31-fold in the spleen, 5,39-fold in the lungs and 3,77-fold in the bone marrow), whereas GFP-3BNC117-treated mice showed minimal reduction across tissues (0,75-fold in the spleen, 1,4-fold in the lungs and 1,73-fold in the bone marrow). CAR-T cell engraftment in tissues was confirmed by CAR or GFP-3BNC117 transgene detection in the spleen and the lung (Fig, 5D).

Circulating 3BNC117scFv-IgG1Fc antibodies were detectable by day 7 post Hybrid CAR administration, with a mean concentration of 135.67 ng/mL (range: 49.04-389.84 ng/mL). By day 14, levels further increased to a mean of 288 ng/mL (range: 57.15-648.78 ng/mL), demonstrating effective and sustained secretion in vivo (Fig. 5E). In GFP-3BNC117-treated mice, 3BNC117scFv-IgG1Fc antibodies were also detected at day 7, reaching a mean concentration 169.8 ng/mL(range: 14.01-548.08 ng/mL). However, levels modestly declined by day 14 to a mean of 128,71 ng/mL (range: 7.68-309.69 ng/mL).

## Discussion

Despite remarkable advances in ART, which have transformed HIV into a manageable chronic condition, a definitive cure remains elusive (55). A sustained cure will require strategies capable of eliminating latent reservoirs responsible for viral rebound upon ART interruption. Numerous curative approaches have been explored, each tackling different aspects of viral persistence, yet facing significant barriers. For instance, the ‘shock and kill’ strategy employs latency-reversing agents (LRAs) to reactivate silent proviruses, rendering infected cells susceptible to immune-mediated clearance. However, clinical studies have demonstrated limited success due to incomplete reactivation, insufficient immune effector responses and HIV immune evasion through MHC-I downregulation (56). Consequently, latency reversal without adequate enhancement of antiviral immunity remains insufficient for durable remission (57–60). In parallel, bNAbs have emerged as promising immunotherapeutics, delaying viral rebound and transiently reducing the reservoir reduction via Fc-effector engagement. Fc-FcγR engagement is now recognized as essential for bNAb efficacy, particularly in driving ADCC and ADCP (61,62). Nonetheless, repeated dosing requirements, viral escape and limited half-life pose challenges to long-term efficacy and feasibility in the context of curative strategies (63–72).

CAR-T cell therapy offers a powerful modality to redirect cellular immunity toward HIV-infected cells in an MHC-independent manner, overcoming an important viral evasion mechanism. A growing body of literature supports the clinical potential of CAR-T cells with next generation approaches enhancing efficacy, durability and resistance to viral escape. For example, Gonda et al. demonstrated that multispecific duoCARs targeting conserved Env epitopes achieved near-complete viral suppression in humanized mice, while Guan et al. introduced cytomegalovirus (CMV)-HIV-specific CAR-T cells, which enable expansion of anti-HIV effector cells following CMV vaccine boosting, potentially addressing the challenge of CAR-T persistence (73–75). Moreover, in vivo engineering strategies are being explored to circumvent ex vivo manufacturing complexity, representing a major step toward scalable deployment and underscoring the translational momentum of this field (76).

However, a critical question remains: can T cells alone eliminate HIV reservoirs and sustain ART-free remission, or must other immune mechanisms be engaged? Given the multifaceted nature of HIV persistence, coupled with the compromised immune responses of PLWH, single-agent treatments are unlikely to sustain ART-free remission (77). Unlike conventional CAR-T cell therapies, the Hybrid CAR platform combines direct cytotoxicity with bNAb secretion, integrating cellular and humoral mechanisms in one product. Previous studies also explored engineering T cells as vehicles for antibody delivery. Powell et al. first demonstrated that HIV-specific T cells secreting bNAbs can elicit ADCC (44). However, MHC-restricted recognition leaves them vulnerable to HIV’s MHC-I downregulation and epitope variation; escape routes circumvented by CAR-T cells. Alternative designs used synthetic Notch (syNotch) receptors to trigger bNAb secretion upon antigen recognition, whereas Hybrid CAR-T cells are expected to constitutively secrete bNAbs, providing continuous output alongside CAR-mediated cytotoxicity (78). More recently, Mao et al. described M10 CAR-T cells, which integrate CAR-mediated killing, bNAb secretion and CXCR5-mediated follicular homing, achieving transient viral load reductions after LRA administration in a Phase I study, although efficacy without an analytical treatment interruption (ATI) remains undefined (79). While this work provides important clinical insights, our study complements it with a thorough preclinical evaluation, including the analysis of Fc-mediated effector function and in vivo activity in a PBMC-humanized mouse model of HIV infection, supporting mechanistic understanding and future clinical translation.

In this study we demonstrate that Hybrid CAR-T cells can combine three complementary activities: (i) CAR-mediated cytotoxicity against HIV-expressing targets, (ii) neutralization of free virions, and (iii) Fc-effector recruitment driving both ADCC and ADCP, integrating cellular and humoral mechanisms in a single product. This design aims to address limitations of endogenous CD8+ T cell responses in HIV, including viral escape, latent reservoirs and MHC-I downregulation which have historically limited viral control (80–83). The CD4-based CAR provides cytotoxicity against Env+ cells, while secreted 3BNC117scFv-IgG1Fc neutralizes free virions and recruits Fc-mediated effectors. CD4-based CARs exploit HIV’s native entry interaction, offering broad, specific recognition via the conserved CD4-binding site and reducing the risk of viral escape, while early generation CD4-based CARs have proven clinically safe (84). Direct comparison with CD4 CAR-T cells revealed that incorporation of the antibody secretion module does not compromise cytotoxic function under standard assay conditions, while providing additional antiviral mechanisms absent from conventional CAR designs. Although Hybrid CAR-T cells retained robust cytotoxic activity comparable to CD4 CAR-T cells, a modest reduction in killing efficiency was observed at lower E:T ratios in a subset of donors. The reduction in cytotoxic activity of Hybrid CAR-T cells at limiting E:T ratios may reflect increased metabolic demands imposed by combining CAR-mediated target killing with antibody synthesis and secretion (85,86). Donor-to-donor variability was observed in transduction efficiency, cytotoxicity, and antibody secretion, mirroring experiences with other adoptive cell therapies and likely reflecting differences in activation state and baseline cytotoxic capacity (87,88). Beyond direct cytotoxicity, Hybrid CAR-T cells exhibited robust functional activation upon antigen encounter. Exposure to HIV Env-expressing target cells induced upregulation of the early activation marker CD69, accompanied by secretion of the effector cytokines TNFα and ΙFNγ. Cytokine production was antigen-dependent, indicating preserved specificity and controlled activation. In parallel, Hybrid CAR-T cells demonstrated antigen-driven expansion in the presence of HIV-infected targets, consistent with productive CAR signalling and proliferative capacity. Despite incorporation of the CD4 extracellular domain, Hybrid CAR expression did not facilitate HIV entry. Together these data indicate that the Hybrid CAR design preserves key features of effective CAR-T cell biology which are critical determinants of in vivo efficacy.

For proof-of-concept, we selected the well-characterized, CD4-binding site bNAb 3BNC117 to be expressed and secreted as an scFv-IgG1Fc fusion protein by the Hybrid CAR-T cells. Although, GFP-3BNC117-transduced T cells secreted higher levels of antibody due to increased transduction efficiency, this did not translate into superior antiviral control. GFP-3BNC117-transduced cells mediated modest reductions in HIV levels in vitro and in vivo, supporting the contribution of antibody-driven antiviral pressure and highlighting the necessity of CAR-mediated cytotoxicity.

To the best of our knowledge, this study provides the first evidence in the context of HIV that CAR-T cell-secreted bNAbs can mediate both ADCC and ADCP, highlighting their dual effector potential. These activities indicate that Hybrid CAR-T cell derived antibodies can effectively engage NK cells and phagocytes, potentially enhancing overall anti-HIV efficacy. Together, these findings highlight a mechanism-integrated approach combining direct cytotoxicity with Fc-effector recruitment that may improve control compared to traditional CAR-T cell therapy alone and warrants validation in models optimized for Fc biology.

For the in vivo exploration of the Hybrid CAR platform, a PBMC-humanized mouse model was employed using PBMCs from HIV-negative donors. Although this model presents limitations, including the risk of GvHD, it is widely used for rapid preclinical assessment (47,89,90). The observed reductions in plasma viremia of Hybrid CAR-T cell-treated mice, coupled with decreased HIV DNA levels across multiple tissue compartments suggest effective viral suppression. Detection of 3BNC117scFv-IgG1Fc in plasma indicates that the engineered T cells can successfully secrete functional antibody in vivo, providing evidence that the cellular platform acts as an endogenous source of 3BNC117, holding the potential for continuous engagement with circulating HIV virions or infected cells. Notably, antibody levels increased between day 7 and day 14 post CAR administration in Hybrid CAR-T cell-treated mice, whereas a moderate decline was observed in GFP-3BNC117-treated controls. This divergence in antibody kinetics could suggest that CAR-mediated activation and expansion of Hybrid CAR-T cells may enhance sustained antibody production and progressive accumulation in vivo. Importantly, the PBMC-humanized model lacks robust reconstitution of innate immune effectors such as NK cells and macrophages, preventing comprehensive evaluation of Fc-mediated mechanisms including ADCC and ADCP in vivo. Therefore, the antiviral effects observed in this system likely underestimate the full therapeutic potential of Hybrid CAR-T cell secreted antibodies in a fully competent immune environment. Detection of CAR-T cell presence in the spleen and the lung supports tissue persistence as a likely contributor to viral suppression. Given the compartmentalized nature of HIV replication, it is reasonable to speculate that local antibody concentrations at sites of active viral activity (e.g., lymph nodes and gut-associated lymphoid tissue) may exceed plasma levels and suffice for physiologically meaningful neutralization and Fc-effector engagement, potentially contributing to a vaccinal effect.

Translating this approach to humans, this strategy could enable sustained antibody delivery without repeated infusions, provided secretion levels, persistence, and safety are established. As part of patient selection, baseline screening for resistance to 3BNC117, along with longitudinal monitoring for the emergence of escape variants post-infusion, should be incorporated into future clinical protocols. Because monotherapy with 3BNC117 may promote the emergence of CD4 binding site escape variants, future iterations of this highly adaptable platform could incorporate complementary bNAbs, such as the V3 glycan-targeting 10-1074 or to be reformatted into bi- or tri-specific formats to broaden epitope coverage and minimize resistance (91).

The ultimate objective of this study is to enable the development of a curative strategy for HIV. By coupling targeted cytotoxicity with sustained in vivo secretion of bNAbs, this system has the potential to reduce treatment burden while enhancing antiviral efficacy through multi-layered immune engagement. Collectively, our findings provide a coherent line of evidence from construct design to preclinical efficacy, supporting Hybrid CAR-T cells as a promising avenue toward functional HIV control.

## Supporting information

Supplementary Material_Broadly neutralizing antibody-secreting CAR-T cells elicit Fc-mediated effector functions in vitro and suppress HIV in humanize

## Data availability statement

All data supporting the findings of this study that are not included in the main text or supplementary materials are available from the corresponding author upon request.

## Author contributions

Conceptualization: W.W., L.V., S.G.

Designed experiments: Z.S., W.W., S.G.

Performed experiments: Z.S., W.W., M.W., E.B., E.D.S., Y.N., M.V.

Analyzed data: Z.S., W.W.

Provided advice: W.W., L.V., S.G., J.V.C.

Writing-original draft: Z.S.

Writing-review and editing: all authors

Visualization: Z.S.

Funding Acquisition: W.W., L.V.

Resources: L.V.

Supervision: W.W., L.V., S.G.

All authors read and approved the manuscript.

