## Supplementary Material_Broadly neutralizing antibody-secreting CAR-T cells elicit Fc-mediated effector functions in vitro and suppress HIV in humanize for "Broadly neutralizing antibody-secreting CAR-T cells elicit Fc-mediated effector functions *in vitro* and suppress HIV in humanized mice"

**LIST OF SUPPLEMENTAL ITEMS**

1. **Supplemental Methods**
2. **Table S1.** Amino Acid sequences of the CD4 CAR, Hybrid CAR and GFP-3BNC117 constructs.
3. **Fig. S1.** Gating strategy for assessment of CD4 CAR and GFP expression on transduced primary CD8+ T cells.
4. **Fig. S2.** Gating strategy for the *in vitro* co-culture assay.
5. **Fig. S3.** Antigen-specific activation of Hybrid CAR-T cells in co-cultures with autologous HIV-infected CD4+ T cells.
6. **Fig. S4.** Antigen-specific expansion of Hybrid CAR-T cells in co-cultures with autologous HIV-infected CD4+ T cells.
7. **Fig. S5.** Hybrid CAR expression on CD4 knock-out cells does not facilitate HIV entry.
8. **Fig. S6.** Raw absorbance values from TZM-bl neutralization assay.
9. **Fig. S7.** Comparison of humanization levels in NSG and NSG-SGM3 mice.
10. **Fig. S8**. Human CD45+ cell engraftment across murine tissues.
11. **Movie S1.** Coculture of NTD cells with autologous HIV-infected CD4+ T cells.
12. **Movie S2.** Coculture of CD4 CAR-T cells with autologous HIV-infected CD4+ T cells.
13. **Movie S3.** Coculture of Hybrid CAR-T cells with autologous HIV-infected CD4+ T cells.

**Table S1.** Amino Acid sequences of the CD4 CAR, Hybrid CAR and GFP-3BNC117 constructs.

| **Construct** | **Amino Acid sequence** |
|---|---|
| CD4CAR | MALPVTALLLPLALLLHAARPGSMNRGVPFRHLL<br>LVLQLALLPAATQGKKVVLGKKGDTVELTCTASQ<br>KKSIQFHWKNSNQIKILGNQGSFLTKGPSKLNDRA<br>DSRRSLWDQGNFPLIIKNLKIEDSDTYICEVEDQKE<br>EVQLLVFGLTANSDTHLLQGQSLTLTLESPPGSSPS<br>VQCRSPRGKNIQGGKTLSVSQLELQDSGTWTCTVL<br>QNQKKVEFKIDIVVLAFQKASSIVYKKEGEQVEFS<br>FPLAFTVEKLTGSGELWWQAERASSSKSWITFDLK<br>NKEVSVKRVTQDPKLQMGKKLPLHLTLPQALPQY<br>AGSGNLTLALEAKTGKLHQEVNLVVMRATQLQK<br>NLTCEVWGPTSPKLMLSLKLENKEAKVSKREKAV<br>WVLNPEAGMWQCLLSDSGQVLLESNIKVLPTWST<br>PVQPSGTTTPAPRPPTPAPTIASQPLSLRPEACRPAA<br>GGAVHTRGLDFACDFWVLVVVGGVLACYSLLVT<br>VAFIIFWVRSKRSRLLHSDYMNMTPRRPGPTRKHY<br>QPYAPPRDFAAYRSKRGRKKLLYIFKQPFMRPVQT<br>TQEEDGCSCRFPEEEEGGCELRVKFSRSADAPAYQ<br>QGQNQLYNELNLGRREEYDVLDKRRGRDPEMGG<br>KPRRKNPQEGLYNELQKDKMAEAYSEIGMKGERR<br>RGKGHDGLYQGLSTATKDTYDALHMQALPPR* |
| CD4CAR-3BNC117scFv-IgG1Fc (Hybrid CAR) | MALPVTALLLPLALLLHAARPGSMNRGVPFRHLL<br>LVLQLALLPAATQGKKVVLGKKGDTVELTCTASQ<br>KKSIQFHWKNSNQIKILGNQGSFLTKGPSKLNDRA<br>DSRRSLWDQGNFPLIIKNLKIEDSDTYICEVEDQKE<br>EVQLLVFGLTANSDTHLLQGQSLTLTLESPPGSSPS<br>VQCRSPRGKNIQGGKTLSVSQLELQDSGTWTCTVL<br>QNQKKVEFKIDIVVLAFQKASSIVYKKEGEQVEFS<br>FPLAFTVEKLTGSGELWWQAERASSSKSWITFDLK<br>NKEVSVKRVTQDPKLQMGKKLPLHLTLPQALPQY<br>AGSGNLTLALEAKTGKLHQEVNLVVMRATQLQK<br>NLTCEVWGPTSPKLMLSLKLENKEAKVSKREKAV<br>WVLNPEAGMWQCLLSDSGQVLLESNIKVLPTWST<br>PVQPSGTTTPAPRPPTPAPTIASQPLSLRPEACRPAA<br>GGAVHTRGLDFACDFWVLVVVGGVLACYSLLVT<br>VAFIIFWVRSKRSRLLHSDYMNMTPRRPGPTRKHY<br>QPYAPPRDFAAYRSKRGRKKLLYIFKQPFMRPVQT<br>TQEEDGCSCRFPEEEEGGCELRVKFSRSADAPAYQ<br>QGQNQLYNELNLGRREEYDVLDKRRGRDPEMGG<br>KPRRKNPQEGLYNELQKDKMAEAYSEIGMKGERR<br>RGKGHDGLYQGLSTATKDTYDALHMQALPPRKE<br>GRGSLLTCGDVEENPGPLEMYRMQLLSCIALSLAL |

| | |
|---|---|
| | VTNSQVQLLQSGAAVTKPGASVRVSCEASGYNIR<br>DYFIHWWRQAPGQGLQWVGWINPKTGQPNNPRQ<br>FQGRVSLTRHASWDFDTFSFYMDLKALRSDDTAV<br>YFCARQRSDYWDFDVWGSGTQVTVSSASTKGPG<br>GGGSGGGGSGGGGSDIQMTQSPSSLSASVGDTVTI<br>TCQANGYLNWYQQRRGKAPKLLIYDGSKLERGVP<br>SRFSGRRWGQEYNLTINNLQPEDIATYFCQVYEFV<br>VPGTRLDLKRTVAAPDKTHTCPPCPAPELLGGPSV<br>FLFPPKPKDTLMISRTPEVTCVVVDVSHEDPEVKF<br>NWYVDGVEVHNAKTKPREEQYNSTYRVVSVLTV<br>LHQDWLNGKEYKCKVSNKALPAPIEKTISKAKGQ<br>PREPQVYTLPPSREEMTKNQVSLTCLVKGFYPSDI<br>AVEWESNGQPENNYKTTPPVLDSDGSFFLYSKLTV<br>DKSRWQQGNVFSCSVMHEALHNHYTQKSLSLSPG<br>K* |
| GFP-3BNC117scFv-IgG1Fc | MVSKGEELFTGVVPILVELDGDVNGHKFSVSGEGE<br>GDATYGKLTLKFICTTGKLPVPWPTLVTTLTYGVQ<br>CFSRYPDHMKQHDFFKSAMPEGYVQERTIFFKDD<br>GNYKTRAEVKFEGDTLVNRIELKGIDFKEDGNILG<br>HKLEYNYNSHNVYIMADKQKNGIKVNFKIRHNIE<br>DGSVQLADHYQQNTPIGDGPVLLPDNHYLSTQSA<br>LSKDPNEKRDHMVLLEFVTAAGITLGMDELYKEG<br>RGSLLTCGDVEENPGPLEMYRMQLLSCIALSLALV<br>TNSQVQLLQSGAAVTKPGASVRVSCEASGYNIRD<br>YFIHWWRQAPGQGLQWVGWINPKTGQPNNPRQF<br>QGRVSLTRHASWDFDTFSFYMDLKALRSDDTAVY<br>FCARQRSDYWDFDVWGSGTQVTVSSASTKGPGG<br>GGSGGGGSGGGGSDIQMTQSPSSLSASVGDTVTIT<br>CQANGYLNWYQQRRGKAPKLLIYDGSKLERGVPS<br>RFSGRRWGQEYNLTINNLQPEDIATYFCQVYEFVV<br>PGTRLDLKRTVAAPDKTHTCPPCPAPELLGGPSVF<br>LFPPKPKDTLMISRTPEVTCVVVDVSHEDPEVKFN<br>WYVDGVEVHNAKTKPREEQYNSTYRVVSVLTVL<br>HQDWLNGKEYKCKVSNKALPAPIEKTISKAKGQP<br>REPQVYTLPPSREEMTKNQVSLTCLVKGFYPSDIA<br>VEWESNGQPENNYKTTPPVLDSDGSFFLYSKLTVD<br>KSRWQQGNVFSCSVMHEALHNHYTQKSLSLSPGK<br>* |

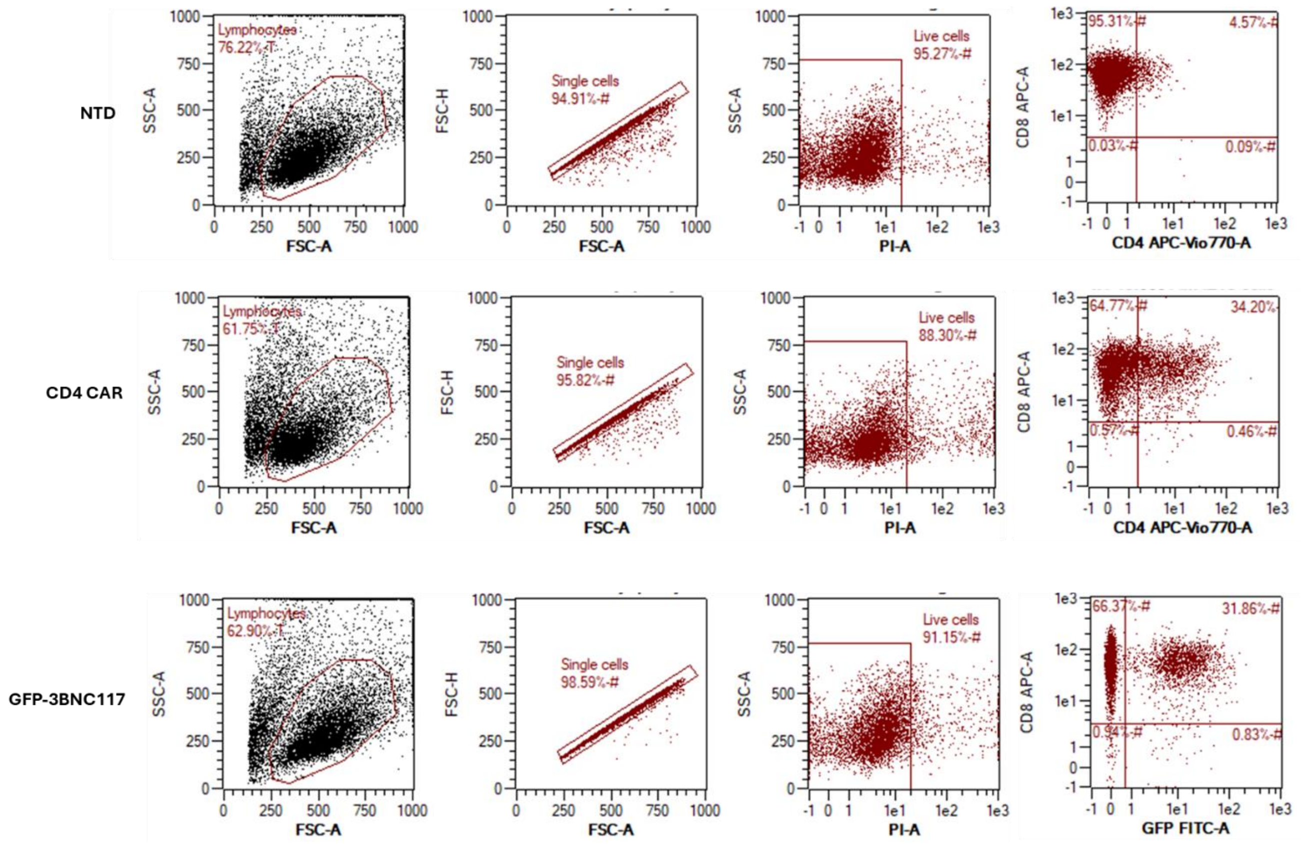


**Supplemental figure 1. Gating strategy for assessment of CD4 CAR and GFP expression on transduced primary CD8+ T cells.**

Primary human CD8+ T cells were initially gated based on forward and side scatter to exclude debris, followed by singlet discrimination. Live cells were identified using a viability dye. Within the CD8+ T cell population, CD4 CAR-transduced cells were identified by CD4 surface expression, whereas GFP-3BNC117-transduced cells were identified based on GFP expression.

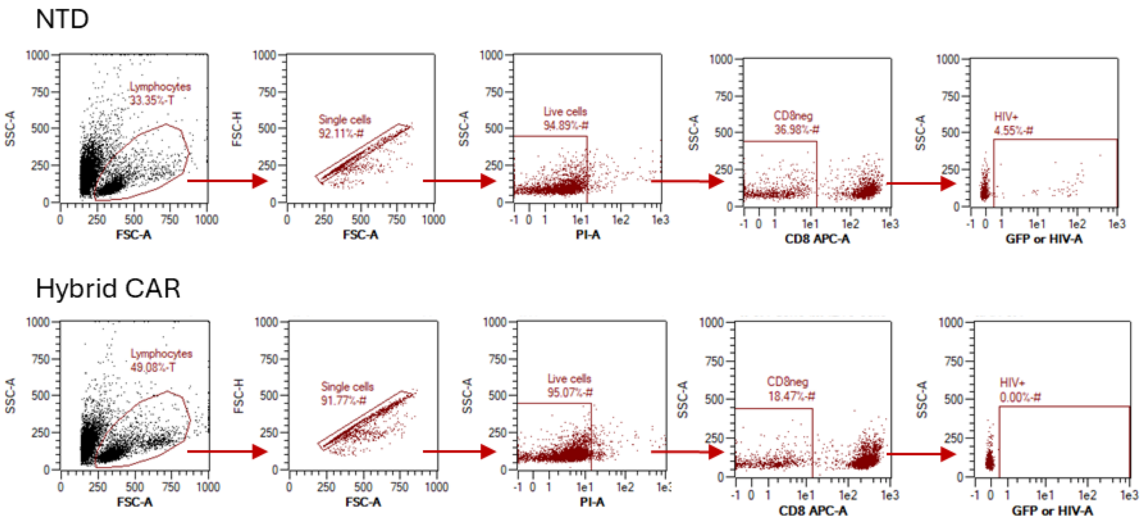


**Supplemental figure 2. Gating strategy for the *in vitro* co-culture assay.**

Gating involves 'single cells' and 'live/dead' cell gating. Subsequently, CD8-positive cells were gated out in order to evaluate percentage of HIV+ cells (marked as GFP+ cells) within the $CD8^{neg}CD4^{neg/pos}$ compartment.

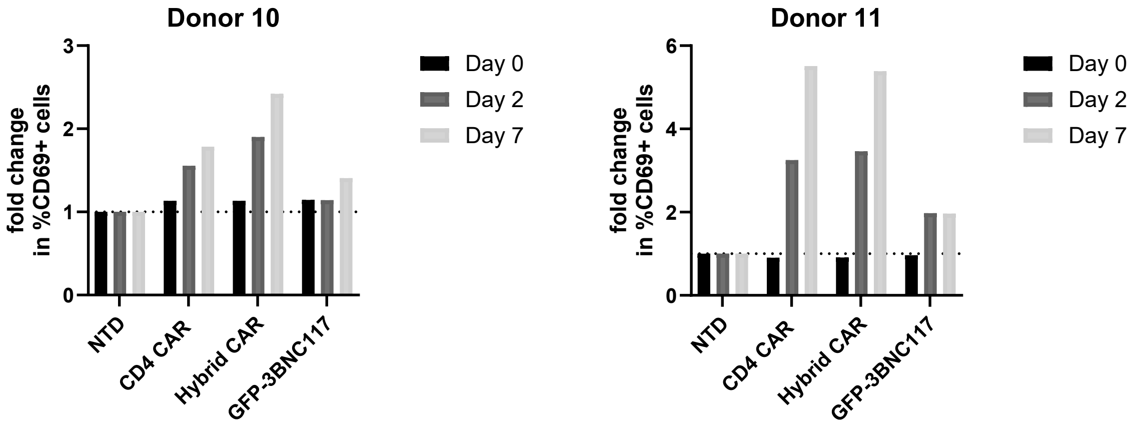


**Supplemental figure 3. Antigen-specific activation of Hybrid CAR-T cells in co-cultures with autologous HIV-infected CD4+ T cells.**

Flow cytometry analysis of (Hybrid) CAR- and GFP-3BNC117-transduced T cells following co-culture with autologous HIV-infected CD4+ T cells. Fold-change analysis of CD69 activation marker expression was performed at Day 0 (pre-co-culture baseline), at Day 2 and 7 of co-culture.

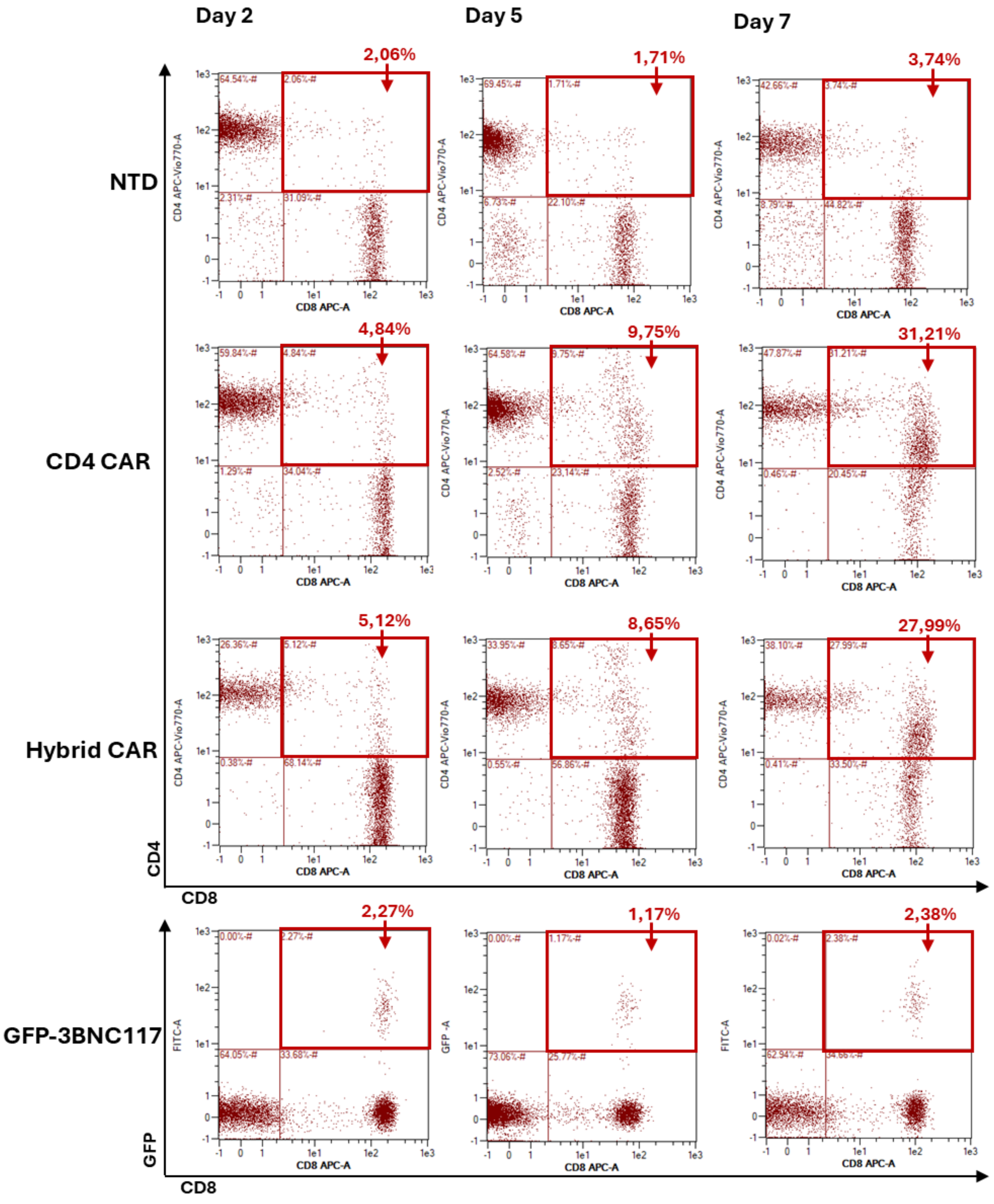


**Supplemental figure 4. Antigen-specific expansion of Hybrid CAR-T cells in co-cultures with autologous HIV-infected CD4+ T cells.**

Flow-cytometry data on Day 2, 5 and 7 of co-culture. (Hybrid) CAR-T cells expand due to target recognition, while levels of GFP-3BNC117-transduced cells remain stable.

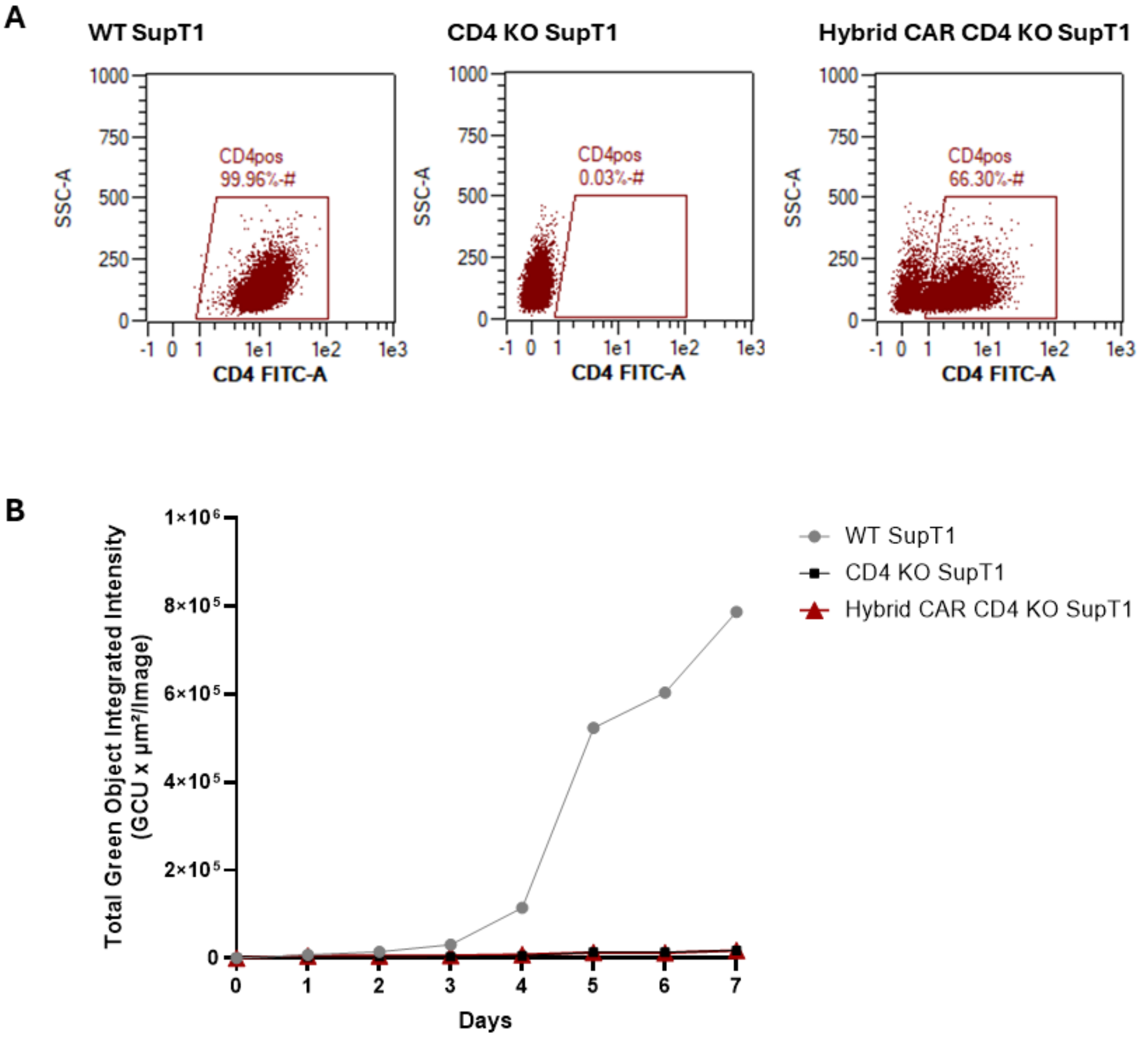


**Supplemental figure 5. Hybrid CAR expression on CD4 knock-out cells does not facilitate HIV entry.**

A. Assessment of CD4 expression by flow cytometry on wild-type CD4+ (WT), on CD4 knock-outs (CD4 KO) and Hybrid CAR-transduced CD4 KO (Hybrid CAR) SupT1 cells. B. Hybrid CAR+ CD4 KO SupT1 cells were exposed to $HIV_{NL4.3\text{-}eGFP}$ and infection was assessed for 7 days, using the IncuCyte S3 Live-Cell Imaging system.

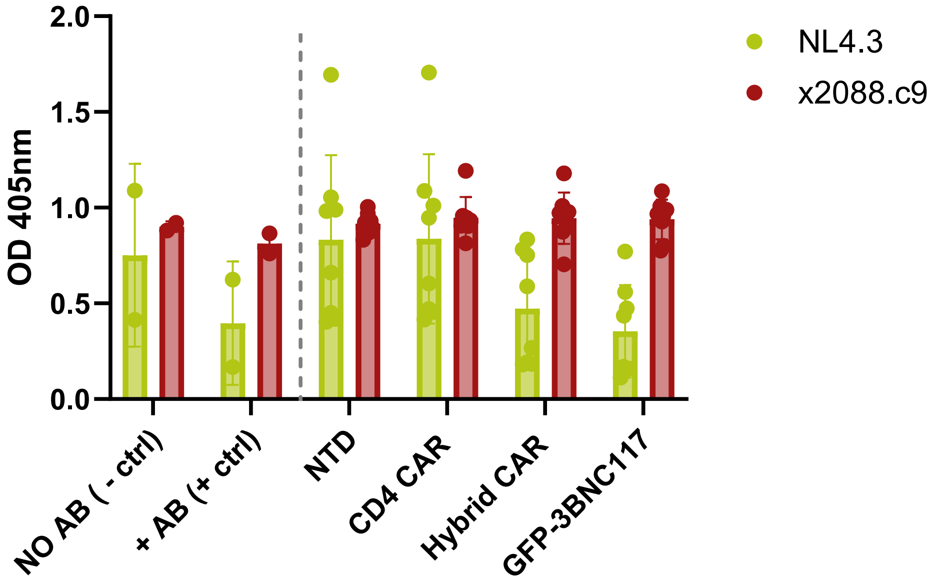


**Supplemental figure 6. Raw absorbance values from TZM-bl neutralization assay.**

Supernatants from non-transduced, (Hybrid) CAR-transduced and GFP-3BNC117-transduced primary T cells were co-incubated with 3BNC117-susceptible ($HIV_{NL4.3}$) or 3BNC117-resistant ($HIV_{x2088.c9}$) strains and applied to TZM-bl cells to evaluate neutralization capacity. Cell culture supernatants can neutralize $HIV_{NL4.3}$ but not $HIV_{x088.c9}$ (Each dot represents one donor; n=8). Abbreviations: -AB (-ctrl), no antibody (negative) control; +AB (+ctrl), recombinant antibody (positive) control; NTD, non-transduced.

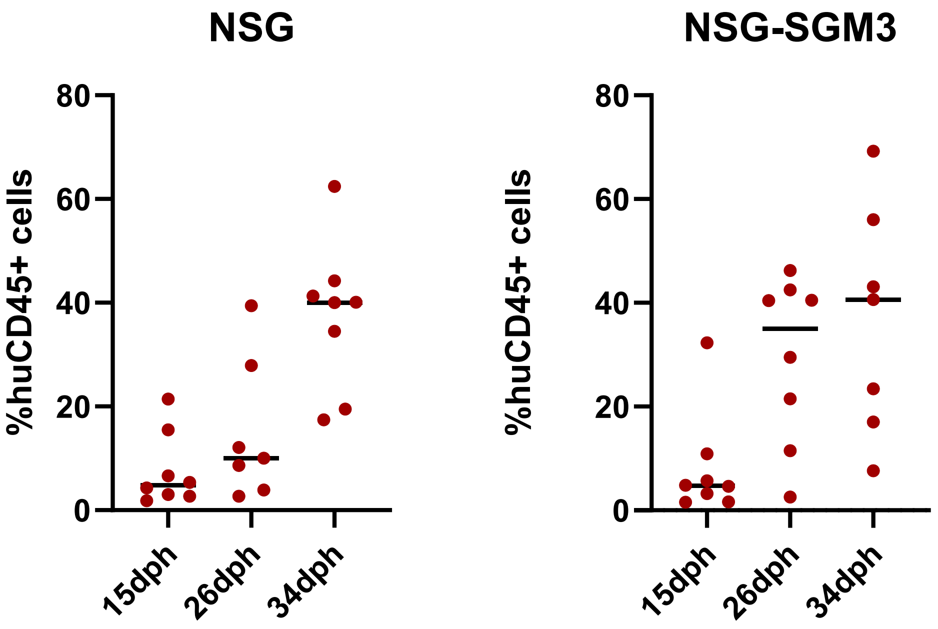


**Supplemental figure 7. Comparison of humanization levels in NSG and NSG-SGM3 mice.**

Levels of human CD45+ cells in peripheral blood were evaluated by flow cytometry at 15, 26 and 34 days post humanization (dph), following retro-orbital intravenous infusion of human PBMCs. NSG-SGM3 mice demonstrated accelerated humanization compared to NSG mice. Each dot represents one animal.

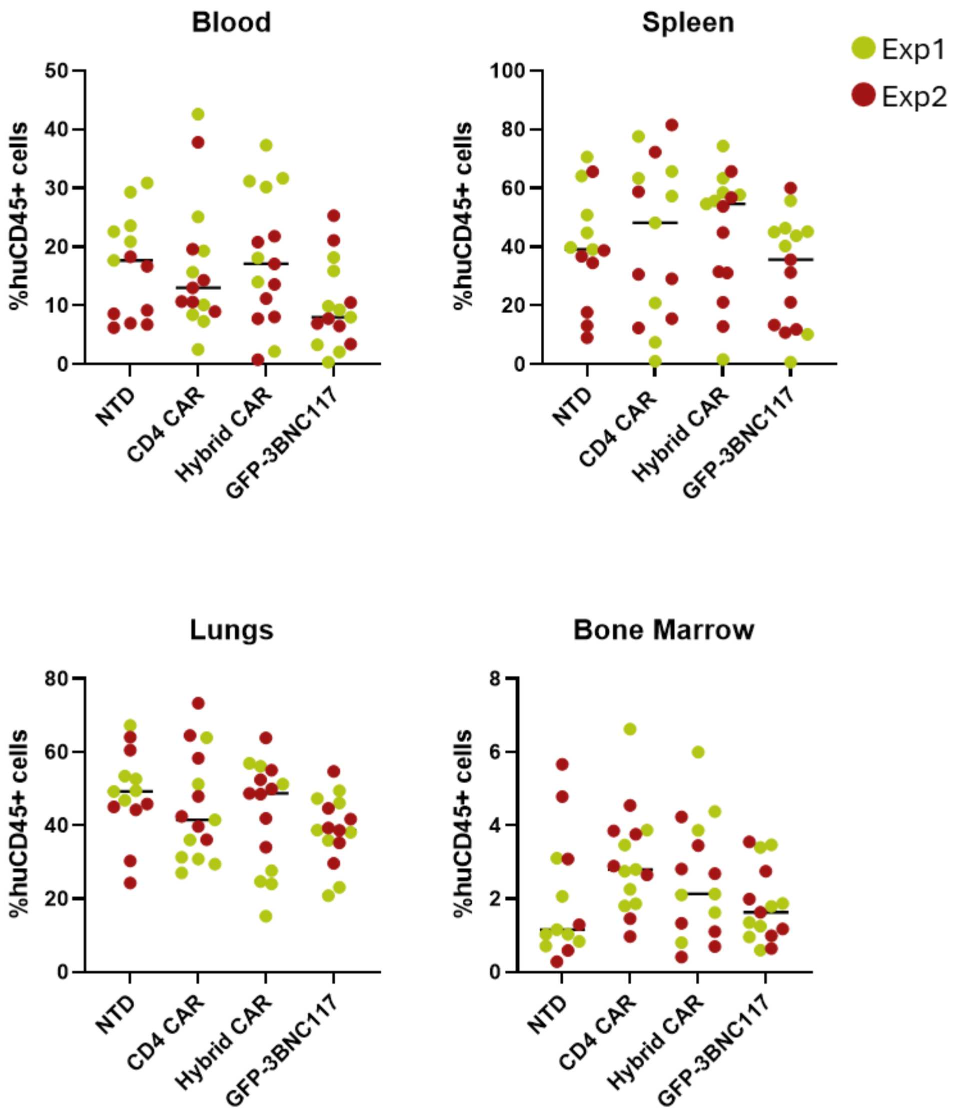


**Supplemental figure 8. Human CD45+ cell engraftment across murine tissues.**

Endpoint analysis of human CD45+ cells in peripheral blood, spleen, lungs and bone marrow. Human cell reconstitution was quantified by flow cytometry.

**Movie S1. Coculture of NTD cells with autologous HIV-infected CD4+ T cells.**

Time-lapse imaging of cocultures between non-transduced (NTD) cells and autologous HIV-infected CD4+ T cells was performed over 7 days, using the IncuCyte Real-time Imaging system.

**Movie S2. Coculture of CD4 CAR-T cells with autologous HIV-infected CD4+ T cells.**

Time-lapse imaging of cocultures between CD4 CAR-transduced cells and autologous HIV-infected CD4+ T cells over 7 days.

**Movie S3. Coculture of Hybrid CAR-T cells with autologous HIV-infected CD4+ T cells.**

Time-lapse imaging of cocultures between Hybrid CAR-transduced cells and autologous HIV-infected CD4+ T cells over 7 days.
